# Pufferfish inflation: drivers of axial and hoop skin strain and regional variation across the body

**DOI:** 10.64898/2026.10.19.760118

**Authors:** Rachel H. S. Tran, Elizabeth L. Brainerd, Samantha M. Gartner, Chiara Micheletti, Andre Vellwock, Bahman Taherkhani, Shahrouz Amini, Ahmad Rafsanjani, Ariel L. Camp

## Abstract

Pufferfish skin undergoes large multi-axial extensions during defensive inflation behaviour, as water is pumped into the stomach and the fish increases its body volume threefold. This remarkable capacity for deformation thus raises questions on how skin extension is spatially distributed across the body, and how skin extension relates to changes in body volume and stomach pressure. However, directly measuring skin extension remains experimentally challenging, particularly when attempting to preserve the body’s complex 3D geometry and natural multiaxial strain during *ex vivo* testing. We used biplanar x-ray videography to measure skin strain (fluoromicrometry) and body volume (dynamic digital endocast) simultaneously with stomach pressure recordings during natural and artificial inflations in porcupinefish (*Diodon holocanthus)*. We found no significant difference in maximum body volumes, body dimensions, peak pressures, and skin extension at maximum inflation between natural and artificial inflation conditions. In all inflations, volume increased steadily while stomach pressure only increased during the last 25-50% of inflation. Skin extension varied biaxially, with mean strains reaching up to 74% in the axial direction and 112% in the hoop direction as the fish became nearly spherical. The greatest skin extensions occurred most dorsocranially (axial) and ventrolaterally (hoop). Strains in the hoop direction were only significantly different between body regions (dorsal, lateral and ventral), with the greatest hoop strains occurring in the ventral body region. Our findings may provide insight into existing hypotheses regarding the relationship between volume, pressure, and skin structure and behaviour, while also providing direct *in vivo* measurements to inform biomimetic design.

**Summary Statement:** Pufferfish skin extends in a complex three-dimensional pattern—with the greatest strains ventrally and in the hoop dimension—as the body inflates into a sphere.

## Introduction

Skin is a vital organ that protects the body and partially determines the limits of body movement and deformation (Kou et al., 2025, Long et al., 1996). The free movement of limbs, flexion of body segments, and bulging of shortening muscles all rely on the skin’s ability to deform three-dimensionally and reversibly. Thus, skin deformation is essential for survival and movement through constantly changing environments (Amini et al., 2025). Yet the skin itself is usually a passive structure with no direct actuation, deforming in response to internal and external forces (Cheney et al., 2022, Yousef et al., 2024).

One extreme example of body and skin deformation is during the inflation behaviour of pufferfishes, such as the porcupinefish *Diodon holocanthus* (**Fig. 1**). In response to predation threats, this species can triple its body volume and erect its dermal spines by pumping water or air into its highly extensible stomach (Brainerd, 1994, Wainwright and Turingan, 1997). This transformation from a smooth torpedo to a spiny sphere is enabled by the pufferfish’s outer surface: a soft-hard interface of hard dermal spines embedded in soft, extensible skin. The skin must undergo remarkable deformation in the form of large multi-axial extensions to achieve body inflation and is thus an extreme example of biological skin mechanics. The kinematics and mechanics of skin extension may also have applications in the bioengineering and biomimetics of synthetic skin-like materials and devices designed to exhibit similar properties of reversible, three-dimensional deformation (Sun et al., 2023, López-González et al., 2023). Nonetheless, there are no direct, multi-axial measurements of skin extension across the body during pufferfish inflation. Nor is it known how skin deformation relates to the increasing body volume and stomach pressure that drives this behaviour. Measuring how the skin deforms in pufferfish would therefore be the first step towards understanding the mechanisms of this remarkably extensible skin, which is also crucial when testing functional morphology hypotheses of skin (Brainerd, 1994, Hertwig et al., 1992) and investigating evolution (Wainwright and Turingan, 1997) and biomimicry of this behaviour. Indeed, pufferfish inflation is already a popular design target for robotic devices (Clarice et al., 2023, Nguyen et al., 2025), even though its kinematics are poorly understood.

**Figure 1.**
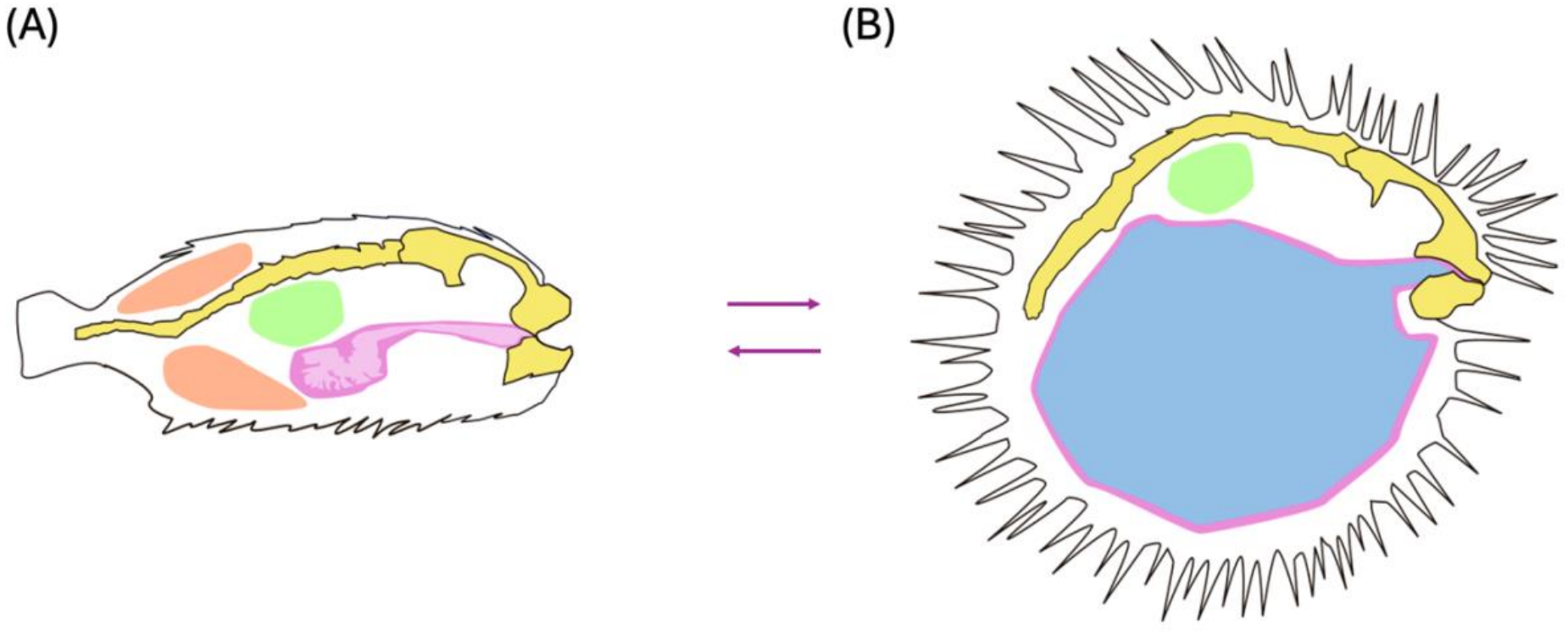
Illustration of the anatomical structures involved in body inflation, based on the porcupine pufferfish *Diodon holocanthus* in a deflated (A) and maximum inflated (B) state. (A, **B)** As the stomach (pink) is filled with water (blue), nearby peritoneum is unfolded (not illustrated), and then the stomach occupies all peritoneal spaces including the potential space (orange) that borders epaxial muscle. **(B)** As maximum inflation is reached and the body becomes fully spherical, the swim bladder (green) is displaced dorsally, and the vertebral column and neurocranium are flexed ventrally (Brainerd, 1994).

Pufferfish skin extension is driven by increasing stomach pressure and volume. Pufferfish use their cranial muscles to pump water from outside, through the buccal cavity and oesophagus, and into the stomach (Wainwright and Turingan, 1997, Wainwright et al., 1995, Brainerd, 1994). As water is pumped in, the volume and pressure of the stomach increases, and it extends into peritoneal spaces. As a result, the surrounding anatomy conforms to increasing stomach volume (**Fig. 1**): muscles stretch, organs and other soft tissues are displaced, the vertebral column flexes, the skin stretches (extends) in all directions, the spines embedded in the skin rotate upwards until they are perpendicular to the skin’s surface, and the volume of the entire body increases. There is no evidence of muscles directly actuating skin extension or spine rotation (Brainerd, 1994), in contrast to, for example, the piloerector muscles used to raise quills in defensive porcupines (Bessa et al., 2024). This paper will not focus on spine motion, so hereafter skin will refer just to the soft tissues of the skin and not these embedded spines.

Instantaneous body volume has not been measured during live pufferfish inflations, although qualitative observations show a gradual increase with each “puff” as water is pumped in (Brainerd, 1994). Meanwhile stomach pressure has been shown to increase non-linearly during live inflations: with pressure remaining low initially and then increasing sharply towards the end of inflation (Brainerd, 1994). The magnitude of skin extensions, temporal and spatial patterns, and relationships to body volume and stomach pressure during pufferfish inflation are unknown. Yet these kinematics can provide insight on the mechanical properties of this hard-soft interface and how such a dramatic body deformation is achieved. For example, Brainerd 1994’s observation of the delayed onset of increasing stomach pressure led to the hypothesis that pufferfish skin has low resistance to extension over most of its deformation range.

Body volume may also increase differently in each body dimension and relative to stomach pressure. For instance, as the body transforms from an initially ellipsoidal (deflated state) to nearly spherical (inflated state) shape, it may expand isometrically: increasing its dimensions at equal rates and magnitudes in the lateral and dorsoventral directions. Or it may expand anisometrically, with different rates in these two directions. We expect craniocaudal dimensions to undergo less change, as this length is largely determined by the flexible, but ultimately incompressible, craniovertebral skeleton. At the level of individual “puffs”, we expect pressure and volume to increase synchronously each time water is pumped into the stomach. But over the entire inflation behaviour, volume and pressure are likely asynchronous (as described in the paragraph above).

Similarly, the magnitude of skin extension may vary in different dimensions and across regions of the fish’s body. Skin may extend equally (isotropically) or unequally (anisotropically) in the longitudinal (axial) and circumferential (hoop) dimensions. While body volume is a single measurement across the entire fish, skin extension may vary across the different regions of the body. Spatial heterogeneity in skin extension would suggest structural properties of the skin and body vary across the fish and in their directionality, which may passively control inflation dynamics. Pufferfish skin extension has only been measured *ex vivo* with uniaxial tensile testing of isolated samples from different species (Brainerd, 1994, Kirti and Khora, 2016). These studies show pufferfish skin has high extensibility relative to other fish (Kennedy et al., 2021, Zhang et al., 2023), with stress (i.e., resistance to extension) only increasing at extensions of approximately 40-60% strain (Brainerd, 1994, Kirti and Khora, 2016). However, these strains have not been directly compared across body regions in the same species. For example, the dorsal skin of *Diodon holocanthus* extended by up to 50% strain in the hoop dimension (Brainerd, 1994), while the dorsal and ventral skin of *Lagocephalus gloveri* only extended by 4-10% strain in the axial direction (Kirti and Khora, 2016). Thus, *ex vivo* skin extension in pufferfish and other fish species suggest that skin extension is unlikely to be homogenous across the body (Madkour et al., 2023, Kenaley et al., 2018, Kennedy et al., 2021).

Now, three-dimensional, x-ray videography methods offer opportunities to measure body dimensions and skin extension in live inflation behaviours, alongside existing pressure recording methods (Brainerd et al., 2010). Fluoromicrometry allows length changes in soft tissues (like the skin) to be measured by tracking the motion of embedded radio-opaque markers with biplanar x-ray video (Camp et al., 2016). Similarly, instantaneous volume can also be measured from biplanar x-ray videos with dynamic digital endocasts (Kaczmarek et al., 2025). These methods can be used during natural, *in vivo* inflations, but also during artificial, indirect actuation of inflation (via a mechanical pump and an *ex vivo* specimen). If such artificial inflations can be shown to accurately replicate *in vivo* patterns, this provides the opportunity for more controlled experiments (with fewer live animals) and would confirm the hypothesis that skin extension is not under direct muscle control.

The aim of this study was to therefore determine the spatial patterns of skin extension and their relationship to pressure and volume during inflation in pufferfish, *Diodon holocanthus.* Biplanar x-ray videography was used to measure skin extension (in axial and hoop directions, at different locations across the body) with fluoromicrometry, as well as to calculate instantaneous body volume over time with a dynamic digital endocast. Stomach pressure throughout inflation was measured simultaneously. Both natural *in vivo* and artificial *ex vivo* inflations were recorded and compared. These data allowed us to determine 1) if body volume increases isometrically and relatively constantly, 2) if skin extension is isotropic or anisotropic, 3) if skin extension is homogenous across the body, and 4) how skin extension is related to body volume and stomach pressure. Our results provide the first direct measurements of skin extension and body volume during inflation, providing insight for existing hypotheses on how volume and pressure are linked to skin structure, and its material properties that previously lacked direct *in vivo* measurements of basic properties. Thus, we provide a better understanding of the biomechanics of the skin and body inflation of pufferfish, as well as its potential applications in biomimetic design.

## Methods

### Animals

Porcupinefish, *Diodon holocanthus* (DH), were purchased from a licenced aquarium supplier and housed in saltwater tanks (24.4–26.2°C and salinity 1.020–1.024) at Brown University. Data in this study were recorded from three individuals DH1, DH2 and DH3 (DH4, DH6 and DH7 in online datasets) with standard lengths of 120 mm, 110 mm and 100 mm and body mass of 135.3 g, 89.8 g and 80.3 g respectively. All procedures were approved by Brown University Institutional Animal Care and Use Committee (24-01-0002) and the University of Liverpool Animal Welfare and Ethical Review Board (AWC0273; AWC0270).

Each fish was surgically implanted with skin and bone markers under anaesthesia (0.09-0.1 g L^-1^ Tricaine Methanesulfonate (MS-222 (Ferndale, Washington, USA)), buffered in seawater) with an analgesic (3 mg kg^-1^ ketoprofen (Schaumburg, Illinois, USA), intramuscular). A total of 38 radio-opaque, 0.5 mm diameter tantalum markers (Brøndby, Copenhagen, Denmark) were implanted in each fish. Four markers were implanted within the neurocranium and upper jaw (which are fused in this species) by incising the skin, drilling 0.5 mm holes directly into the underlying bone, and press-fitting a marker into each hole. The remaining 34 markers were injected into the skin via a hypodermic needle to form an evenly distributed grid, unilaterally over the right side of the fish and avoiding anatomical landmarks that could not be implanted i.e., the eye and pectoral fin (**Fig. 2A**). Note that one marker, the dorsal most caudal (tail) marker was missing in DH1. The skin of *D. holocanthus* has a thick (approximately 2-4 mm in these individuals) layer of dermis. Markers were injected just below the epidermis to lodge in the thick dermis, with the 0.5 mm diameter marker size not expected to affect the mechanical behaviour of the skin. Data collection only occurred after fish resumed normal swimming and feeding behaviour (no more than 24-48 hours).

**Figure 2.**
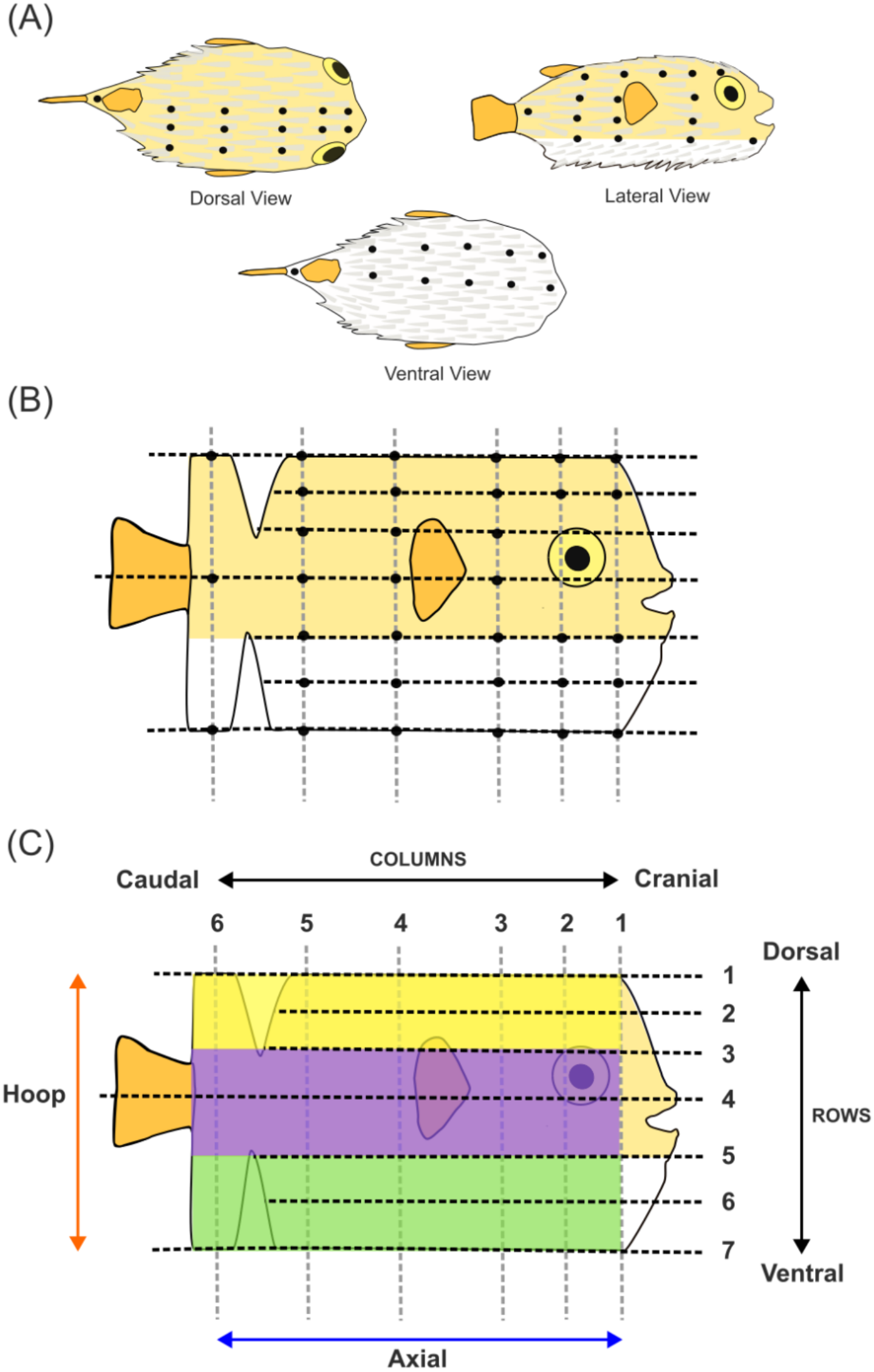
Diagram of skin marker grid for measuring body dimensions, body volume and skin strain. (**A**) Skin markers (black circles) were implanted over the dorsal, lateral, and ventral surface of the fish’s right-side in an approximate grid pattern. (**B**) This diagram depicts the skin of the right side, from the mid-dorsal to mid-ventral line, spread out to show the full area in a 2D lateral view. Markers were placed at the intercepts of dorsoventral columns (vertical grey dashed lines) and craniocaudal rows (horizonal black dashed lines). This formed an approximate mesh, except where interrupted by the eye, jaw, pectoral fin, and tapering of the body towards the caudal fin. (**C**) Skin strain in the axial (cranial-caudal) direction was measured between adjacent markers in the same row (blue arrow). Skin strain in the hoop (dorsal-ventral) direction was measured between adjacent markers in the same column (orange arrow). Strain was compared across three body regions: dorsal (yellow; rows 1-3), lateral (purple; rows 3-5), and ventral (green, rows 5-7).

### Data Collection

Biplanar x-ray videos of inflation were recorded from two dorsal-oblique views (**Fig. 3**) using a custom biplanar x-ray system (Imaging Systems and Services, Painesville, OH, USA) with Flare (12M180MCX, IO Industries, London, Ontario, Canada) high-speed cameras at Brown University. X-ray videos were recorded at 50 frames s^-1^, 2000 μs exposure, and 2048 × 2048-pixel resolution using x-ray energies of 73-90 kV and 200 mA. All inflations were recorded in an acrylic aquarium with 17.6 cm depth of salt water, within a recording volume of ∼ 14.5 (W) × 14.5 (L) × 10 (D) cm. Standard grid and calibration objects were used to remove distortion introduced by x-ray machines and to calibrate the three-dimensional (3D) space (Brainerd et al., 2010).

**Figure 3.**
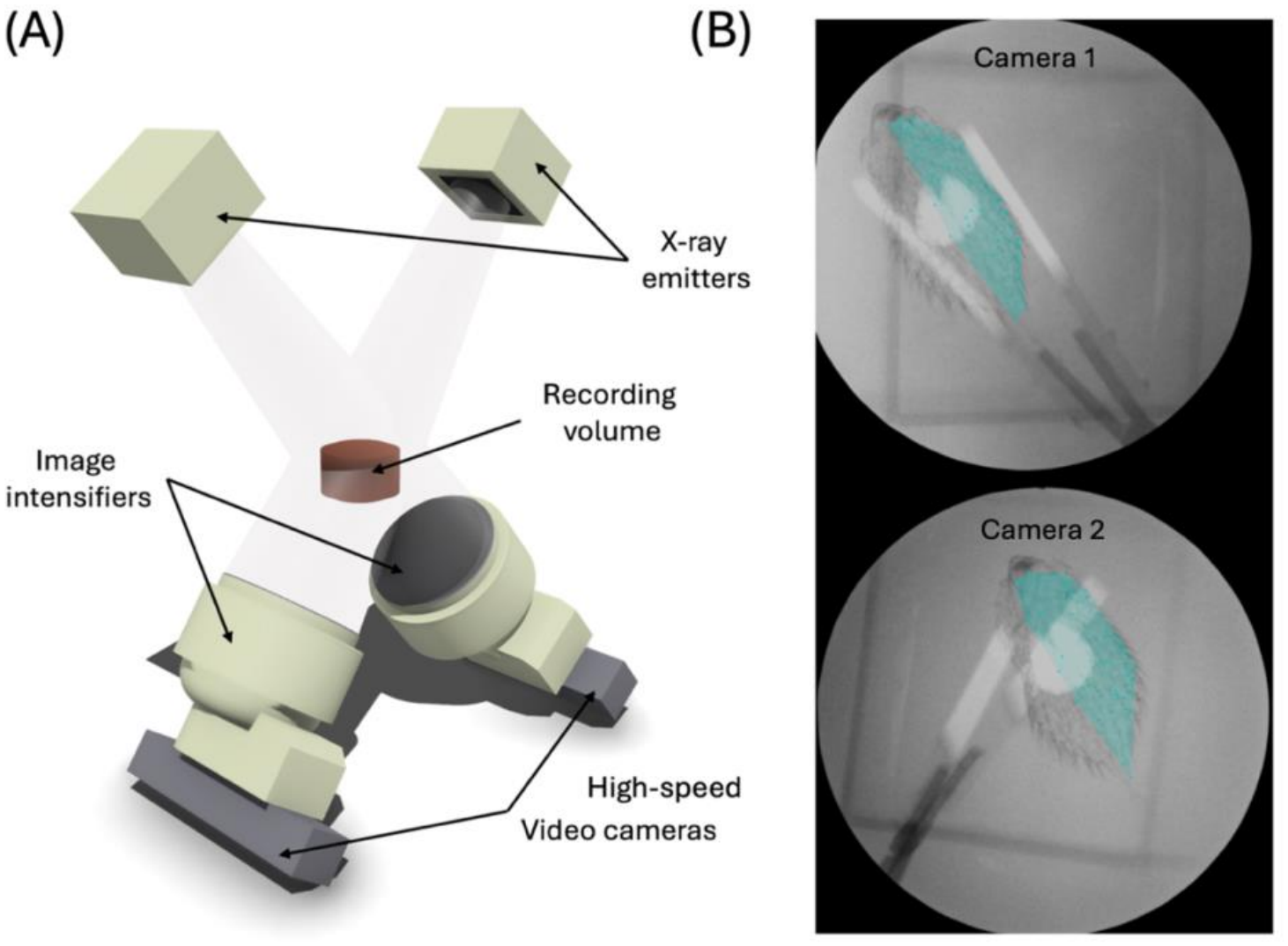
(A) Bi-planar x-ray system setup and (B) example x-ray output images overlaid with a dynamic endocast. (**A**) Labelled 3D rendering of the biplanar x-ray setup for pufferfish inflation recordings. X-ray emitters and their corresponding image intensifiers and high-speed video cameras were positioned obliquely to shoot through water. (**B**) Dorsal-oblique views of the same individual in both x-ray camera views with an endocast fitted around the neurocranium and skin markers on the right side of the fish to estimate body volume. The 3D setup rendering was generated in Shapr3D, UK v5.840.0.

Stomach pressure recordings were synchronously recorded alongside x-ray videos via a SPR-407 pressure transducer (Millar, Houston, Texas, USA) inserted through a flexible polyethylene cannula and recorded at 1000 Hz with a PowerLab 16/35 DAQ system, Bridge Amp and LabChart v8.8.1 (ADInstruments, Central Dunedin, Dunedin, New Zealand). Pressure probe insertion involved lightly anesthetising the fish (0.9g L^-1^ MS-222 in seawater) before threading the cannulated pressure probe through the operculum, out through the mouth, then turning it to pass through the oesophagus and into the stomach. The cannulated pressure probe was then sutured in position at the base of the pectoral fin with 4-0 nylon. Pressure measurements were calibrated daily by recording the voltage as the probe was moved through a 10 cm change in saltwater depth.

We recorded *in vivo* (natural) inflations and post-mortem *ex vivo* (artificial) inflations from each fish. Food was withheld from the fish one day prior to filming ensure an empty stomach and avoid regurgitation (Brainerd, 1994). Natural inflations were initiated by applying bilateral pressure to the fish’s hyomandibula, via a long-handled wood and plastic tongs, to mimic a predation threat. Compression was maintained until the end of inflation: cessation of buccal pumping and a plateau in stomach pressure. Fish were given 10-15 minutes between inflation trials to avoid fatigue (McGee and Clark, 2014). In both natural and artificial inflations, it was necessary to stabilise the fish against the bottom of the aquarium to position the tongs at the start of the inflation. The handler then lifted the fish slightly off the bottom for most of the recorded event.

Due to the risk of fish biting through the pressure probe and a faulty probe, stomach pressure was recorded successfully during natural inflations in only one fish (DH3). After natural inflation recordings, fish were euthanised by an overdose (2 g L^-1^) of buffered MS-222 in salt water. Immediately after death was confirmed, a cuffed 4.0 endotracheal tube (Dynarex, Montvale, New Jersey, USA) was inserted through the mouth and into the stomach and the cuff inflated to create a watertight seal within the oesophagus (**Fig. S1**). The external opening of the endotracheal tube was connected to an aquarium pump (MAXI-JET 600, MarineLand, Blacksburg, Virginia, USA). The specimen was immersed in the same volume and depth of salt water as used for natural inflations, and the pump turned on to deliver water into the specimen’s stomach while biplanar x-ray video and pressure were recorded. The pump was left to run until maximum inflation was reached, at which point the endotracheal tube was often ejected from the oesophagus. Pressure was recorded for all artificial inflations. In total, 3, 4 and 5 useable natural inflation recordings were obtained from DH1, DH2 and DH3 respectively. Meanwhile 2, 4 and 2 artificial water inflation recordings were obtained from DH1, DH2 and DH3 respectively.

Computed tomography (CT) scans were taken post-mortem of each fish at 1632 × 1112-pixel resolution and 0.041 mm slice thickness with a SkyScan 1276, Bruker scanner and reconstructed in NRecon (Bruker, Billerica, Massachusetts, USA). Scans were segmented in 3DSlicer v5.6.2 ((Kikinis et al., 2014); https://www.slicer.org/) to create polygonal mesh models of the markers and neurocranium. Models were imported into Autodesk Maya 2024 (San Rafael, California, USA) and the xyz (3D) coordinates of each neurocranium marker calculated with custom scripts from the ‘XROMM MayaTools’ package by David Baier, Steve Gatesy, and Armita Manafzadeh (https://bitbucket.org/xromm/xromm_mayatools).

Raw data (x-ray videos, calibration and grid images; CT scans; pressure recordings) are publicly available and stored on the XMAPortal (https://xma.xromm.brown.edu/) in the Public Dataset ‘Skin Strain in Pufferfish Inflation’. Video data are stored with their essential metadata in accordance with best practices for video data management in organismal biology (Brainerd et al., 2017).

### Body Volume

The 3D positions of all markers were tracked using XMALab v2.2.5 ((Knörlein et al., 2016), software and instructions available at https://github.com/XROMMPackages/xmalab) with a minimum tracking precision of 0.1 mm across all co-osseus marker pairs. See the publicly available data for tracked 3D coordinates ([FigShare](https://figshare.com/)) of implanted skin markers throughout a sample of inflation trials. Rigid body transformations were calculated for the neurocranium to create marker-based XROMM animation of this bone. 3D coordinates of all markers (excluding the two left-most neurocranium markers: left eye and left jaw) were used to calculate changes in body volume using a dynamic digital endocast (Kaczmarek et al., 2025). Marker coordinates were loaded into MATLAB vR2023b (The MathWorks Inc, Natick, Massachusetts, USA) to generate alpha shapes and calculate volumes at each frame (alpha setting 5). These were then imported into Autodesk Maya to visually confirm that the body endocast aligned with the x-ray video images and XROMM-animated neurocranium. Because skin markers were placed unilaterally, endocast volume was doubled to calculate total body volume. Raw estimates of body volume were then filtered (10^th^ order low pass Butterworth filter with a 0.3 half-power frequency and zero-phase digital filtering), and peak volume and local maxima and minima extracted using a custom MATLAB script (**Fig. 4A**).

**Figure 4.**
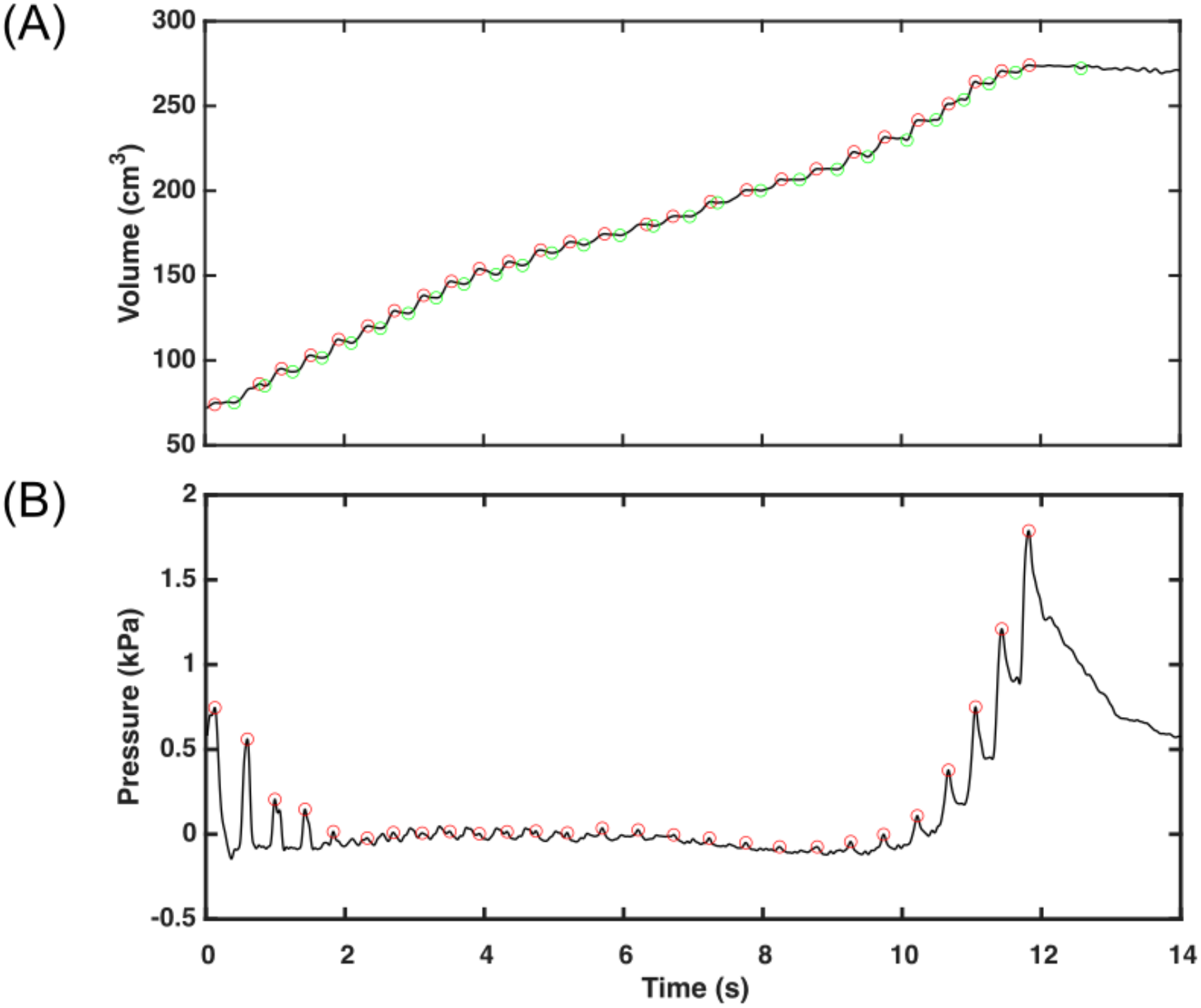
Identification of local maxima of (A) body volume and (B) stomach pressure. Red circles highlight local maxima, and green circles (A only) highlight local minima with each ‘puff’ taken during a natural inflation trial.

### Pressure

Stomach pressure was filtered (10^th^ order low pass Butterworth filter with a 0.03 half-power frequency and zero-phase digital filtering) and baseline corrected before extracting peak pressures, and local maxima and minima using a custom MATLAB script (**Fig. 4B**).

### Skin Strain and Body Dimensions

Skin strain was calculated over time as the change in 3D vector distance between adjacent skin markers from the x-ray videos using fluoromicrometry (Camp et al., 2016). All strain calculations were done in custom MATLAB scripts using the same 3D marker coordinates as the dynamic endocast (excluding any neurocranium markers). The identity of each skin marker was confirmed, before strains were calculated as the change in intermarker distance (relative to initial) divided by the initial intermarker distance between specified marker pairs. Initial distance was measured at the first frame of each trial, prior to the start of inflation. Axial strain was calculated along a craniocaudal axis between adjacent pairs of markers within the same row (**Fig. 2B, 2C**). Hoop strain was calculated along a dorsoventral axis between adjacent pairs of markers within the same column (**Fig. 2B, 2C**). For example, axial strain was calculated between markers located where row 1 intercepts columns 1 and 2 (markers R01C01 and R01C02). While hoop strain was calculated between markers located where column 1 intercepts rows 1 and 2 (markers R01C01 and R02C01). Strains in both directions were also grouped according to body region (**Fig. 2C**): dorsal (rows 1-2), lateral (rows 3-4), and ventral (rows 5-6 for hoop and rows 5-7 for axial strain). For each fish, we additionally calculated the mean axial and hoop strain independently for each marker pair for both inflation conditions. For the analysis at maximum inflation (defined as peak body volume), strains from all trials were averaged across marker pairs but individuals kept separately when looking at differences between body regions.

The 3D coordinates of skin markers were also used to calculate overall body dimensions. Dorsoventral height was defined as the distance between markers R01C03 and R07C03, craniocaudal length as the distance between R07C01 and R07C06, and mediolateral width as twice the distance between R04C04 to the midpoint of R01C04 and R07C04. Both skin strain and body dimensions were filtered using the same method as body volume, with a 10^th^ order low pass Butterworth filter with a 0.3 half-power frequency and zero-phase digital filtering.

### Statistical Analysis

To determine whether inflation condition (natural or artificial) and individual influenced maximum body volume, body dimensions, peak stomach pressure, and skin strain, data were first assessed for normality using the Shapiro–Wilk test. Variables were then analysed using a linear mixed-effects ANOVA with Satterthwaite approximation. Where a significant effect was detected, additional post hoc analyses were performed using a modified linear mixed-effects model to assess differences between live and artificial measurements within each individual. Pairwise comparisons between individuals were then conducted using Holm–Bonferroni correction while averaging across inflation conditions.

To compare strain types (axial and hoop) independently against body region (dorsal, lateral and ventral), we conducted a linear mixed-effect ANOVA with Satterthwaite approximation to evaluate overall differences. Where significant, a post-hoc test for pairwise comparisons between body regions was also performed with *p*-values adjusted using the Holm–Bonferroni correction method.

Finally, to compare axial and hoop strains within the same body region in all individuals a Wilcox rank sum test was performed and *r* values calculated. All figure generation and statistical analysis was facilitated by custom MATLAB scripts.

## Results

### Inflation Behaviour

We measured a total of 20 inflations (12 natural and 8 artificial) from three pufferfish. Body volume, body dimensions, and skin strain were calculated for all 20 inflations. Due to technical limitations, pressure was only recorded for 13 of the 20 inflations (5 natural, 8 artificial). The variables (e.g., strain, volume and pressure) from natural inflations showed a stair-step like pattern over time, reflecting the individual ‘puffs’ as the pufferfish pumped water into their stomachs (**Fig. 4**). This pattern was absent in artificial inflations, where a mechanical pump provided a constant flow of water into the stomach.

No significant difference in the magnitudes of body volume, stomach pressure or skin strain at maximum inflation was observed between inflation conditions or individuals (**Table 1**), with the exception of body dimensions. Craniocaudal length and mediolateral width had a small but significant difference between individuals and inflation conditions (see ‘Body Volume and Shape’ section below and **Table S1**). We therefore report mean ± standard deviation values pooled across all individual and inflation conditions unless stated otherwise.

**Table 1.** Mean values ± standard deviation and linear mixed-effects ANOVA results between inflation conditions, individuals, body volume, body dimensions and stomach pressure at maximum inflation.

|  | Individual | *N* Trials<br>(nat, art) | Mean Value<br>Natural | Mean Value<br>Artificial | *F*<br>(df1, df2) | *p* |
|---|---|---|---|---|---|---|
| Max Body Volume (cm<sup>3</sup>) | DH1 | (3, 2) | 250.03 ± 20.57 | 256.16 ± 9.19 | 0.0002<br>(1,17) | 0.988 |
|  | DH2 | (4, 4) | 190.84 ± 25.14 | 191.95 ± 22.69 |  |  |
|  | DH3 | (5, 2) | 203 ± 8.73 | 196.86 ± 18.99 |  |  |
| Dorsoventral height (mm) | DH1 | (3, 2) | 87.31 ± 0.23 | 87.32 ± 0.07 | 0.002<br>(1,17) | 0.965 |
|  | DH2 | (4, 4) | 81.39 ± 0.28 | 80.85 ± 0.34 |  |  |
|  | DH3 | (5, 2) | 78.09 ± 0.55 | 78.81 ± 0.02 |  |  |
| Craniocaudal length (mm) | DH1 | (3, 2) | 98.825 ± 0.258 | 98.76 ± 0.1 | 13.6<br>(1,17) | 0.002 |
|  | DH2 | (4, 4) | 81.93 ± 0.43 | 83.18 ± 0.27 |  |  |
|  | DH3 | (5, 2) | 85.44 ± 0.77 | 86.71 ± 0.13 |  |  |
| Mediolateral width (mm) | DH1 | (3, 2) | 82.81 ± 0.16 | 83.06 ± 0.15 | 15.72<br>(1,17) | 0.001 |
|  | DH2 | (4, 4) | 76.67 ± 0.28 | 77.28 ± 0.1 |  |  |
|  | DH3 | (5, 2) | 83.22 ± 0.041 | 83.68 ± 0.07 |  |  |
| Peak Pressure (kPa) | DH1 | (-, 2) | - | 2.36 ± 0.17 | 0.38<br>(1,13) | 0.551 |
|  | DH2 | (-, 4) | - | 2.87 ± 1.64 |  |  |
|  | DH3 | (5, 4) | 2.22 ± 1.16 | 2.23 ± 0.15 |  |  |
| Axial Strain | DH1 | (3, 2) | 0.3 ± 0.21 | 0.38 ± 0.2 | 1.607<br>(1,160) | 0.207 |
|  | DH2 | (4, 4) | 0.29 ± 0.26 | 0.29 ± 0.25 |  |  |
|  | DH3 | (5, 2) | 0.33 ± 0.25 | 0.4 ± 0.25 |  |  |
| Hoop Strain | DH1 | (3, 2) | 0.47 ± 0.3 | 0.82 ± 0.34 | 0.003<br>(1,144) | 0.959 |
|  | DH2 | (4, 4) | 0.5 ± 0.41 | 0.48 ± 0.38 |  |  |
|  | DH3 | (5, 2) | 0.55 ± 0.38 | 0.52 ± 0.33 |  |  |

Natural (nat) and artificial (art) inflation trials. Degrees of freedom (df).

### Body Volume and Shape

During inflation pufferfish increased their body volume by a mean of 2.66 ± 0.65 times, and up to 3.5 times, their initial body volume. Under both inflation conditions, volume increased at an approximately linear rate until reaching and maintaining a maximum value (**Fig. 5A, 5B**). Maximum inflation was defined as the time point where maximum body volume occurred. During natural inflations, pufferfish pumped water into their stomachs with a mean total of 28.6 ± 5.5 puffs over 9.27 ± 2.46 seconds to fully inflate. Each puff increased body volume by a mean of 7.37 ± 3.47 cm^3^, with a body volume flow rate of 22.3 ± 36.34 cm^3^ s^-1^ (**Fig. 5A**). For many puffs, local volume maxima were also followed by a local minima indicating a loss of body volume (2.52 ± 2.92 cm^3^) (**Fig. 4A**). In *ex vivo* artificial inflations, body volume increased slightly faster at 23.2 ± 13.41 cm^3^ s^-1^ over 9 ± 2.43 seconds to fully inflate; **Fig. 5B**).

**Figure 5.**
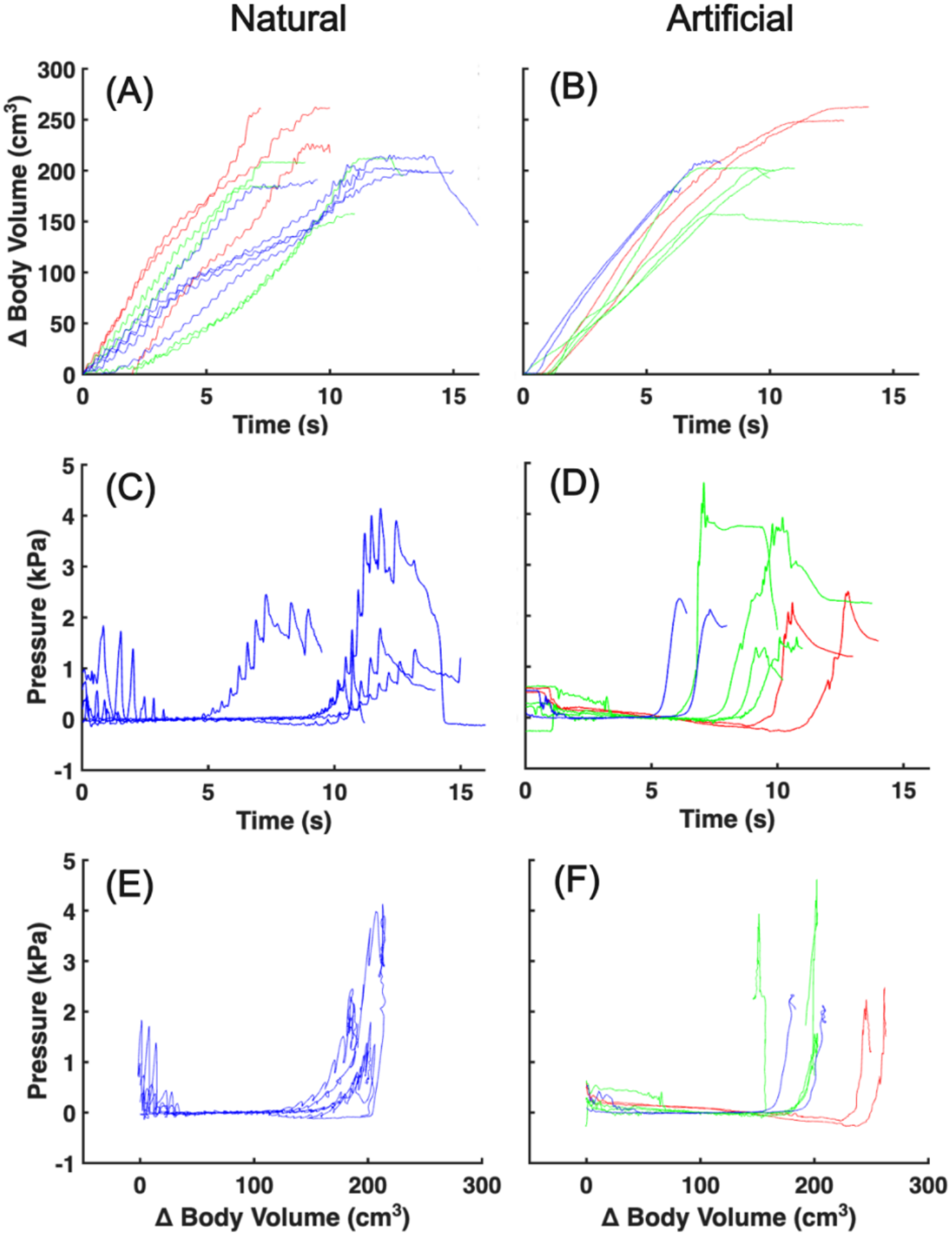
Change in body volume and stomach pressure during inflation. Body volume and stomach pressure of natural (**A**, **C**, **E**) and artificial (**B**, **D**, **F**) inflations where red is DH1 (3 natural and 2 artificial trials), green is DH2 (4 natural and 4 artificial trials) and blue is DH3 (5 natural and 2 artificial trials). Stomach pressure in natural inflations was measured only in DH3, due to technical limitations.

Pufferfish also changed shape from approximately ellipsoidal shape when deflated to nearly spherical at full inflation (**Table 1**; **Fig. 6**). Before inflation, the craniocaudal length of each fish was about twice the dorsoventral height and mediolateral width (**Fig. 6**). During inflation, mediolateral width and dorsoventral height approximately doubled, while craniocaudal length decreased by about 10%, so that at maximum inflation all three dimensions were nearly equal as required for a sphere (**Fig. 6**).

**Figure 6.**
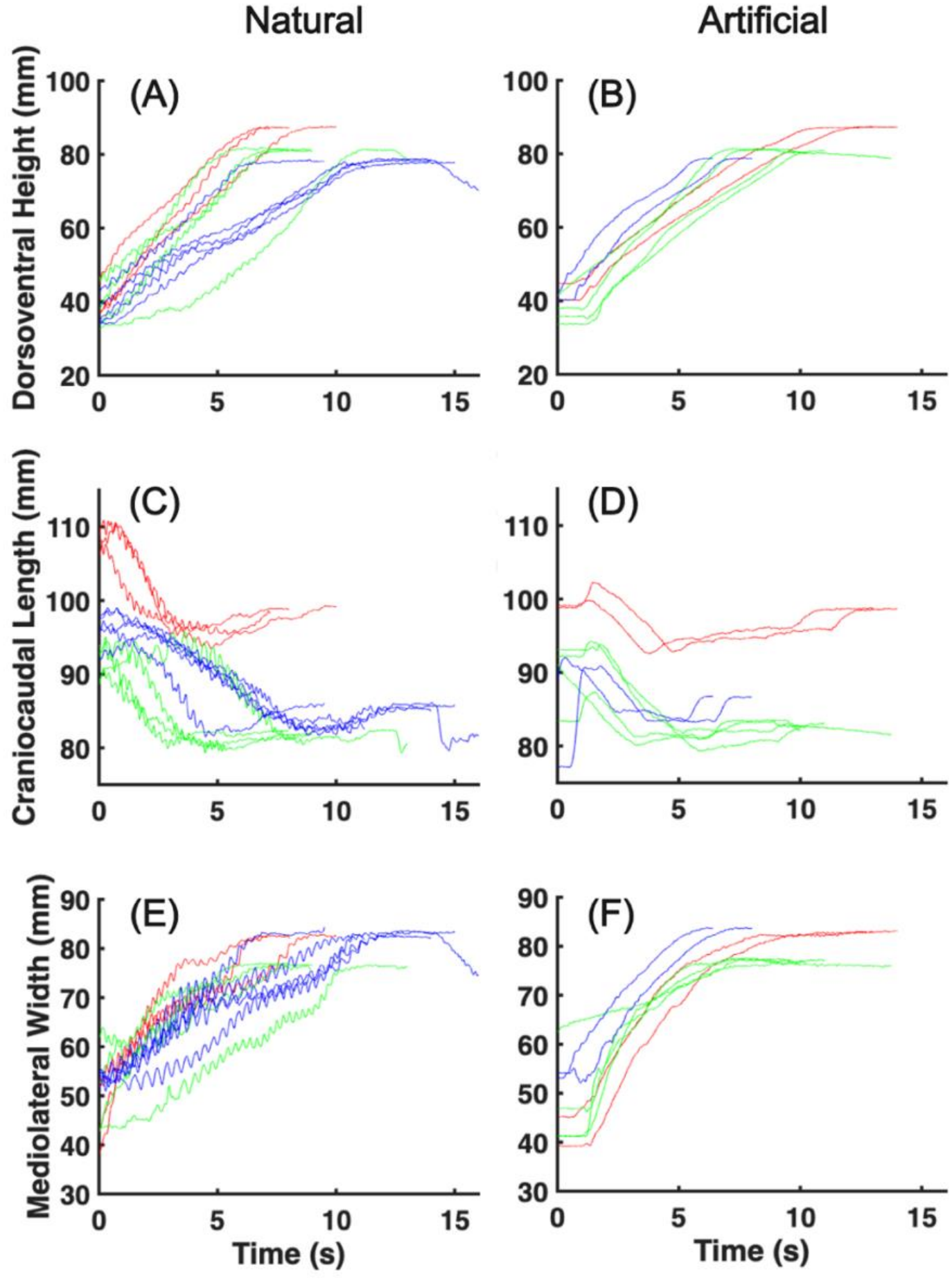
Body dimensions of all natural and artificial inflation trials. (**A, B**) Dorsoventral height, (**C, D**) craniocaudal length and (**E, F**) mediolateral width. Column 1 is natural inflation trials (*n* = 12) and column 2 is artificial inflation trials (*n* = 8). In all subplots red is DH1 (3 natural and 2 artificial trials), green is DH2 (4 natural and 4 artificial trials) and blue is DH3 (5 natural and 2 artificial trials).

There was a small but significant difference in the craniocaudal length and mediolateral width at maximum inflation between individuals and inflation conditions (**Table 1**). Average craniocaudal length at maximum inflation was approximately equal across inflations for DH1, but 1 mm greater in artificial inflations for DH2 and DH3 (**Table 1**). Average mediolateral width was consistently greater in artificial inflations by up to about 1 mm (**Table 1**). Additional investigations between natural and artificial trial measurements of craniocaudal length and mediolateral width revealed no significant differences within individuals (all *p*>0.05; e.g., DH1 natural vs. DH1 artificial craniocaudal lengths; **Table S1**). But significant differences were observed between individuals (all *p* < 0.05; e.g., all DH1 craniocaudal lengths vs. all DH2 craniocaudal lengths; **Table S1**).

### Stomach Pressure

Stomach pressure increased relative to initial values by 1.37–4.61 kPa (*n* = 13, peak pressure 2.45 ± 1.10 kPa), over 2 phases. First, in the ‘stable’ phase pressure remained almost constant for the first 50–75% of inflation (**Fig. 5C, 5D**). In natural inflations, there were cycles of troughs and peaks with each puff during this stable phase, but the peaks of increasing pressure were relatively small (−0.023 ± 0.143 kPa change per puff) and balanced by nearly equal troughs of decreasing pressure (**Fig. 5C**). In artificial inflations, pressure remained low and relatively constant during the stable phase, with some downward baseline drift, likely due to changes in hydrostatic pressure as the handler moved the fish up slightly in the water column (**Fig. 5D**). Second, in the ‘exponential’ phase of natural inflations, pressure increased rapidly by at least 0.05 kPa per puff with a mean of 0.31 ± 0.24 kPa per puff during this phase (**Fig. 5C**). In artificial inflations, pressure increased rapidly by 1.51 ± 3.54 kPa s^-1^ without any cycles (**Fig. 5D**). All inflations had a stable and exponential phase of pressure and the transition between them occurred once body volume had increased by about 100-150 cm^3^ (approximately double the deflated body volume) (**Fig. 5E, 5F**). In some natural and artificial inflations there was a brief period of pressure increase early in the inflation process due to the handler stabilising the fish against the bottom of the aquarium to position it correctly before raising it up into the water column (**Fig. 5C, 5D**).

### Skin Strain

For simplicity, we only quantitatively report skin strain at a single time point: maximum inflation, i.e., the time of peak body volume. Strains are reported relative to initial length, with positive values denoting an increase in length (extension). Skin extension was almost always observed at maximum inflation, with strains averaged across all local marker pairs of 0.32 ± 0.16 and 0.51 ± 0.30 in the axial and hoop direction, respectively (**Fig. 7** in summary and **Figs. S2, S3** for all trials). Rarely, negative (shortening) strains were measured, generally in the axial direction and in one of the most cranial (row 2, between column 1-2) or caudal (row 1, between column 5-6) locations (**Figs. S2, S3**).

**Figure 7.**
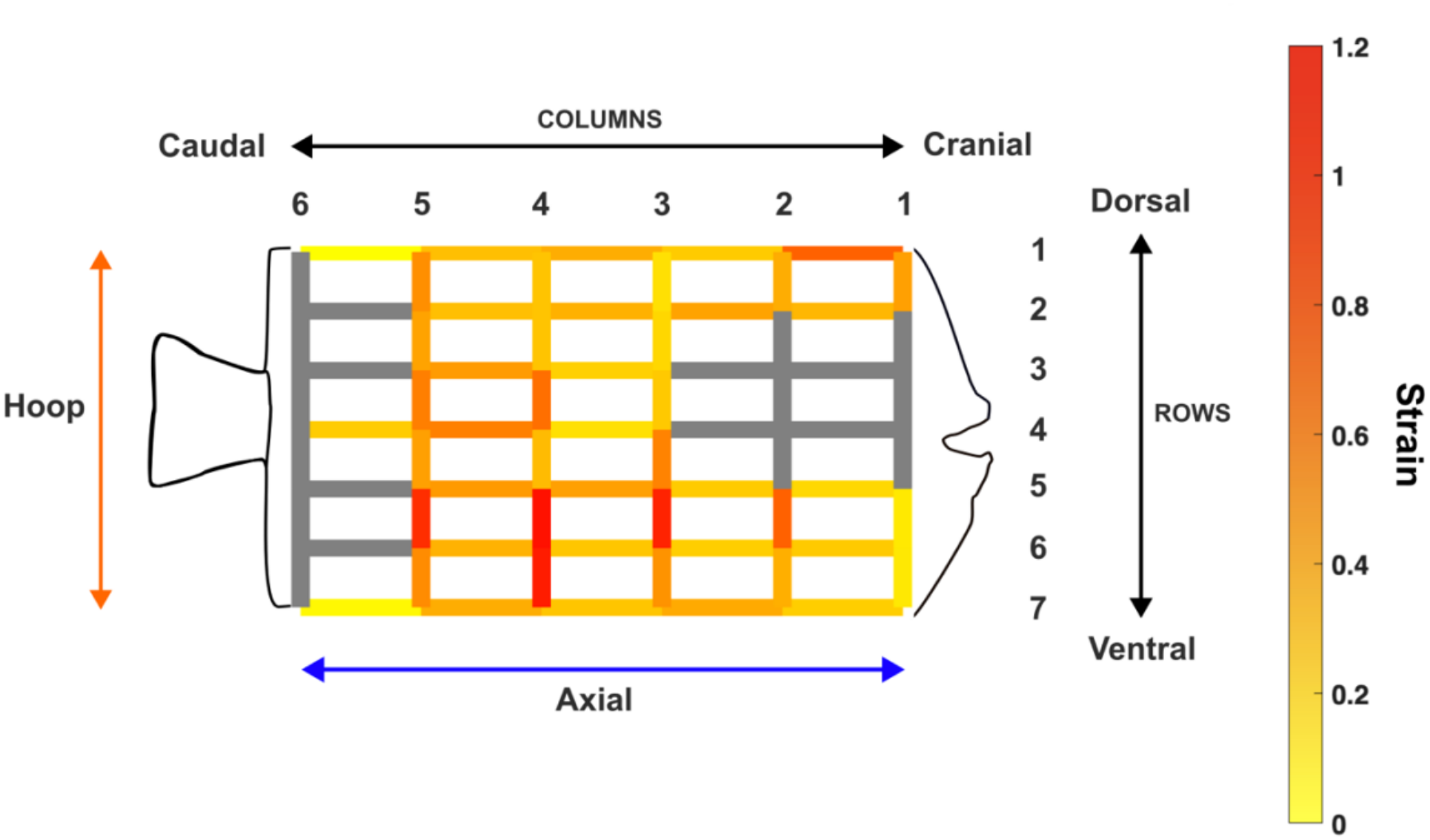
Colourmap of axial and hoop skin strain across the body. Mean axial (rows, blue arrow) and hoop (columns, orange arrow) skin strain across all individuals and inflation trials (*n* = 20 trials from 3 individuals). The colour bar (right) indicates strain magnitudes. Grey lines are locations where no skin strain was calculated due to anatomical landmark obstruction e.g., the eye and pectoral fin.

At the level of local skin strains between adjacent markers, we observed a wide range of mean axial (0.01-0.74) and hoop (0.10-1.12) strain magnitudes at maximum inflation (**Fig. 7**). The greatest local axial strain occurred in a single cranial, dorsal, nearly mid-sagittal location: across rows 1-2 and column 1 (0.74 ± 0.45). In contrast, the greatest local hoop magnitudes occurred in four ventral, caudal, lateral (near the border of white and pigmented skin locations across rows 5-6 and columns 2-5 (0.97 ± 0.16) (**Figs. 2, 7**). Again, the highest local strains occurred in the hoop direction (**Fig. 7**). Skin strain variation within and among individuals was large, with standard deviations ranging from 0.07-0.50 for axial strain and 0.07-0.55 for hoop strain. This variability is illustrated on a trial-by-trial basis in **Figs. S2, S3**.

At the level of body regions (dorsal, lateral, ventral), skin strain magnitudes also varied widely but only differed significantly across the body in the hoop direction when pooling inflation conditions. Axial skin strain did not differ significantly amongst body regions (LME, F_2,80_ = 0.517, *p* = 0.599), with a mean value of 0.33 (± 0.03) across all regions (**Fig. 8A**). Meanwhile, values of hoop skin strain did differ significantly amongst body regions (LME, F_2,72_= 6.49, *p* = 0.003) (**Fig. 8B**), with ventral strains (mean 0.65 ± 0.46) being greater than dorsal (mean 0.34 ± 0.17) strains (*p_Holm* = 0.002). Hoop strain was only significantly greater than axial strain in the lateral (*p* = 0.045, *z* = −2.01, *r* = −0.35) and ventral *p* = 0.0013, *z* = −3.26, *r* = −0.39) regions. Dorsal region strains did not differ between axial and hoop directions (*p* = 0.961, *z* = 0.049, *r* = 0.0069).

**Figure 8.**
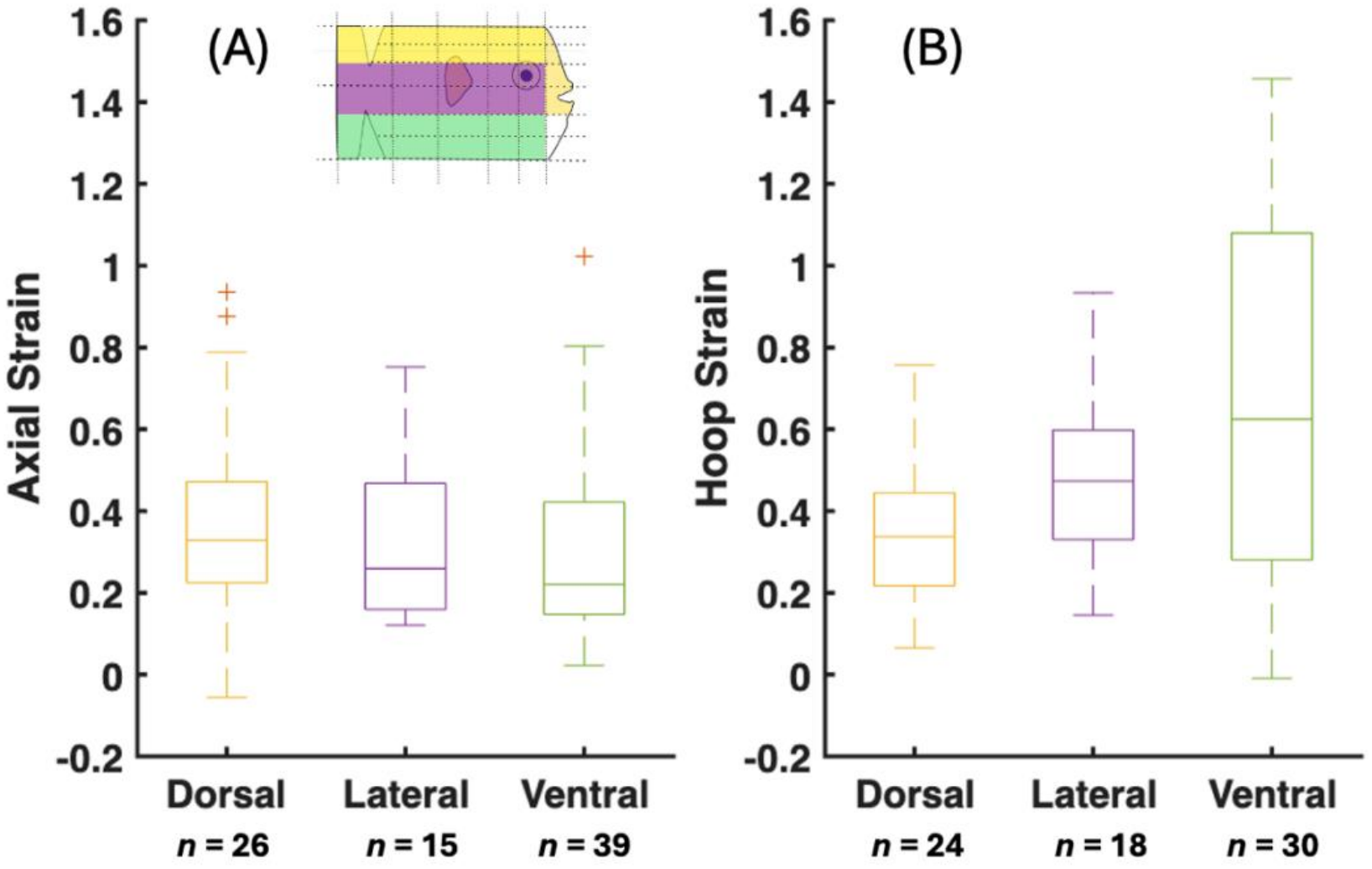
Skin strains across different body regions. (**A**) Axial and (**B**) hoop strains grouped by body region (dorsal, lateral, ventral), where all trials were averaged for each fish. Skin strains from the dorsal body region are yellow, lateral is purple and ventral is green. N indicates the number of marker pairs in each group from 3 fish. The top, middle and bottom of each box is the 75^th^ percentile, median and 25^th^ percentile respectively. The whiskers extending above and below each box are maximum and minimums. Red plus signs are outliers.

## Discussion

We recorded the first direct measurements of three-dimensional body volume and skin extension, as a function of time, during the inflation behaviour of the porcupine pufferfish, *Diodon holocanthus*. The skin of this pufferfish underwent large extensions with mean strains of up to 74% and 112% in the axial and hoop direction respectively. Skin extension was anisotropic and heterogeneous across the body, with the largest skin extensions concentrated dorsocranially (axial) and ventrolaterally (hoop). In contrast, body volume increased approximately isometrically in the mediolateral and dorsoventral directions, suggesting skin structure and/or material properties may be altering the relationship between body volume and skin extension. Most skin extension and body volume increase occurred while stomach pressure was low, indicating that the skin requires little force to extend. Together these results show how the skin of pufferfish enables such remarkable 3D deformation and generates new hypotheses of its structure-function relationships and biomimetic potential.

### Body Volume and Stomach Pressure

Body volume increased relatively constantly during inflation, while stomach pressure only increased during the last 25-50% of inflation (i.e., the ‘exponential phase’). This confirmed the delay between the onset of volume increase and onset of pressure increase previously proposed by Brainerd (1994). Most body volume increase occurred during the ‘stable phase’ of low stomach pressure, suggesting the body is highly compliant and extensible over this range of volumes. The initial peaks (maxima/minima) of pressure recorded in some trials appear to be artefacts. For *in vivo* trials, these were likely caused by the initial positioning the fish at the bottom of the tank, towards the centre of the recording volume, and placing the tongs on the subjects hyomandibula used to mimic predation threat. For *ex vivo* trials, these were likely the result of motion of the entire fish when the mechanical pump was turned on.

Body volume increased through nearly equal increases in mediolateral width and dorsoventral height. As expected craniocaudal length was relatively constant, decreasing by <10% from deflated to inflated state. Observations of the x-ray videos show this craniocaudal shortening is due to ventral flexion of the intervertebral and craniovertebral joints at maximum inflation, as proposed by Brainerd, 1994. Thus, the craniovertebral skeleton itself was not compressed, but as it flexed the straight-line distance between the rostrum and the tail decreased.

Peak body volume and pressure values were similar in natural and artificial inflations, showing that the mechanical pump and euthanised specimen successfully replicated maximum body inflations. This supports the hypothesis that the magnitude and shape of inflation are driven by stomach inflation and material properties of the fish, not direct muscular control of the skin. The rate of volume increase was greater in artificial inflations, as the mechanical pump generated a higher flow rate than the buccal pump. However, we did not observe any evidence of over-inflation in artificial inflations, suggesting live pufferfish may already be inflating near their maximum mechanical capacity. Peak volume magnitudes were similar to Brainerd,1994 at about three times resting body volume. This suggests our dynamic digital endocast method is reasonably accurate, despite not fully capturing surface curvature (which would lead to slight underestimation). It should also be noted that these volumes reflect the entire body, not just the stomach. As the volume of the skeleton and soft tissues should remain constant, we expect our body volume rates to be a good representation of the rate of stomach volume increase and its change relative to initial volume.

Peak stomach pressures, however, were all lower than the 8 kPa (60 mmHg) reported by Brainerd, 1994. This difference may be owed to the positioning of the pressure transducer within the stomach and/or the motivation of the fish (natural) or tightness of the seal of the mechanical pump (artificial). If the latter two, we may be underestimating peak pressure performance.

During natural inflations, we recorded the cyclical stomach pressure and body volume changes generated by the buccal pump. We observed the same step-like pattern in pressure found previously (Brainerd, 1994), and for the first time show a similar pattern in volume. With each puff, pressure and volume increased and then decreased slightly. This slight decrease in body volume may be an artefact of water moving within the body that is not fully captured by the endocast (e.g., towards the tail). This interpretation is consistent with the decreasing magnitude of local minima (i.e., less water loss) towards maximum inflation, as the skin becomes stiffer and limits further redistribution of water. On the other hand, decreases in stomach pressure were more pronounced. The changes in stomach pressure reflect the forces required to actively expand the skin and stomach. The small peaks and troughs observed during the early phase of inflation may be explained by inertia associated with the mass of tissue being displaced, even when the skin is not yet resisting expansion due to stiffness. Meanwhile, towards maximum inflation, skin extension is increasingly resisted by skin stiffness, resulting in progressively larger increases in stomach pressure with each puff. The larger decreases in pressure following these spikes are likely attributable primarily to relaxation of the skin after stretching. This stress-relaxation behaviour is characteristic of viscoelastic tissues such as skin (Clancy et al., 2010). We therefore hypothesise that the inertial force required to expand the skin remains approximately constant across volumes, and that the larger pressure decreases observed later in inflation are therefore predominantly associated with viscous relaxation.

### Skin Extension during Inflation

Our measurements showed that pufferfish skin is exceptionally extensible relative to previous *ex vivo* measurements made in pufferfish and other fish species. Locally, pufferfish skin extended by means up to 74% and 112% of its initial length in the axial and hoop direction respectively at maximum inflation. To our knowledge, these are the highest skin extension strains measured in fish. Hoop strains observed in the ventral body region of this study were similar to the 40% hoop strain (140% extension from rest) previously measured from *Diodon holocanthus* with isolated, *ex vivo* samples taken from the abdominal/ventral body region (Brainerd, 1994). Outside of pufferfishes, the closest comparable measurements of fish skin strain are of *Eptatretus stoutii* (Pacific hagfish), that experienced peak stresses between ∼ 19-25 MPa (1.9 × 10^7^ – 2.5 × 10^7^ Pa) at ∼ 35-45% strain under uniaxial testing in both the axial and hoop direction (Kennedy et al., 2021). The strain measurements in this study were calculated from the straight-line distances of adjacent markers, which do not account for body contouring or skin curvature. Thus, we may be slightly underestimating skin strain.

Skin extension at maximum inflation followed an anisotropic pattern in hoop versus axial strain and was heterogeneous across the body. The highest local strains were all in the hoop dimension (**Fig. 7**) and lateral and ventral regional strains were significantly greater in the hoop than axial dimension (**Fig. 8**). Regionally, only hoop strains differed significantly between the dorsal, lateral, and ventral regions. However, strains within each region were highly variable and local strains show variation across the body. For axial strain, the greatest extension occurred in a single craniodorsal region (**Fig. 7**, row 1, column 1). This region is near where the skin attaches to the rigid structure of the upper jaw. The difference in mechanical properties and constraint between the rigid jaw and the more compliant skin may result in a non-uniform distribution of deformation, with strain becoming concentrated near this transition. Consequently, the craniodorsal region exhibited the greatest axial extension. For hoop extension, the greatest extension occurred in a ventro-lateral group of regions (**Fig. 7**, rows 5-6, columns 3-5). This area is near the widest part of the fish when deflated (ellipsoidal), does not require flexion of the craniovertebral column to expand fully (like the dorsal region), has fewer underlying organs (like the ventral region) and is likely to follow circumferential expansion of the body when inflated.

We hypothesise that the variation in skin extension magnitudes across the body and directions reflects both the body geometry of the fish and the microstructure of the skin. First, the pufferfish body does not maintain a uniform shape but transforms from an approximately elliptical deflated fish to a nearly spherical inflated fish. The deformation—and therefore skin extension—required for this geometrical transformation will vary across the body. Second, microstructure of the skin may result in more compliance—and therefore more extension—in the hoop dimension. The skin of *Diodon holcanthus* has a collapsed/compact collagen layer arrangement that requires relatively little force to stretch in the hoop dimension (Brainerd, 1994). Axial strain and biaxial strain in response to known forces have not been measured in this species. We hypothesise that the skin architecture allows greater hoop than axial strain for a given force. Skin architecture and material properties have also been shown to vary between dorsal and ventral body regions in other pufferfish species (Madkour et al., 2023, Kirti and Khora, 2016). We therefore also hypothesise that the regional differences in *Diodon* may also reflect variation in skin architecture. However, investigations into both geometry and skin architecture and material properties have yet to be done.

## Conclusion

We provide the first direct measurements of multiaxial skin strain during the inflation of live pufferfish. Skin strain varies with direction and body region, and the resultant increase in body volume occurs primarily during low stomach pressure. Our results show not only exceptional magnitudes of skin extension compared to other fish, but also a complex three-dimensional pattern that varies across the body, in two dimensions, and throughout time. We hypothesise that the skin deformation of pufferfish is actuated by increasing stomach pressure and volume, but it is also shaped by the geometry of the fish’s body and the architecture of the skin itself. Further research into these areas will improve our understanding of how such extreme skin and body deformation is achieved and controlled, and how it can be mimicked artificially.

## Acknowledgements

We are grateful to Erika Tavaras for support with experiments and CT scanning at Brown University, and Peter Falkingham for assistance with CT analysis. Chiara Micheletti acknowledges funding support from the Alexander von Humboldt Foundation, and from the Health Engineering, Nano, and Materials Science Areas of Advance at Chalmers University of Technology.

## Author Contributions

Rachel H. S. Tran (RT), Elizabeth L. Brainerd (EB), Samantha M. Gartner (SG), Shahrouz Amini (SA), Chiara Micheletti (CM), Andre Vellwock (AV), Ahmad Rafsanjani (AR), Bahman Taherkhani (BT), and Ariel L. Camp (AC).

Conceptualisation (All)

Methodology (RT, AC, EB, SG)

Formal analysis (RT)

Investigation/Data Collection (RT, AC, EB, SG)

Data curation (RT)

Writing - original draft (RT, AC)

Writing – review & editing (All)

Visualisation (RT, AC)

Supervision (AC)

Project administration (AC)

Funding acquisition (AC, SA, AR)

## Data Availability

Raw data (x-ray videos, pressure recordings, CT scans) deposited in XMAPortal (Study ‘Skin Strain in Pufferfish Inflation’, https://xma.xromm.brown.edu/) will be available on publication.

Processed data (marker xyz coordinates, stomach pressures, body volumes) will be available through FigShare on publication.

