## Supplementary Table and Figures for "Pufferfish inflation: drivers of axial and hoop skin strain and regional variation across the body"

### Supplementary Materials

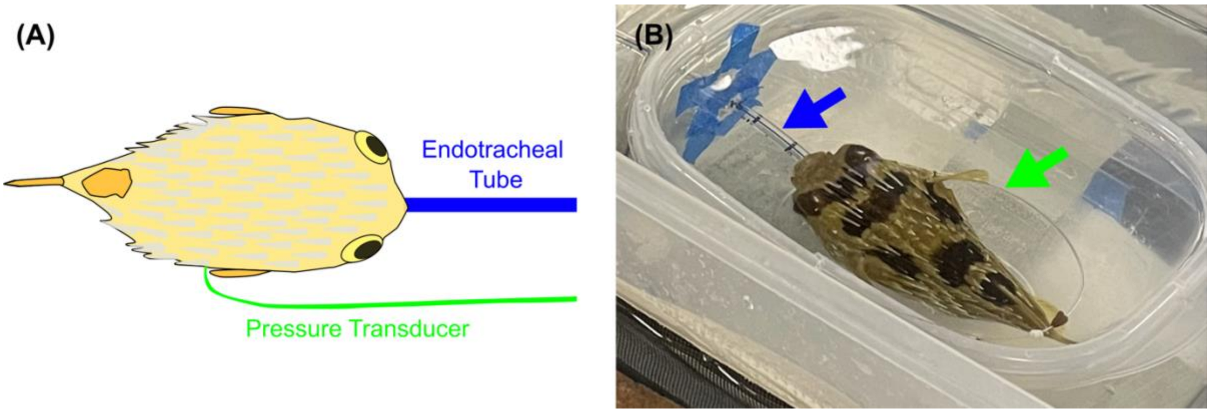


**Figure S1. Experimental setup for artificial inflation trials.** (**A**) Illustration of the setup used for artificial inflation. A cuffed endotracheal tube (blue) was inserted into the specimen's oesophagus, and then its cuff inflated to create a watertight seal. The tube was then connected to a water pump that delivered salt water into the specimen's stomach. A pressure transducer (green) was secured at the base of a pectoral fin. (**B**) Photograph of the artificial inflation setup, showing the endotracheal tube (blue arrow) and pressure transducer (green arrow).

**Table S1. Linear mixed-effects ANOVA and Holm–Bonferroni post hoc test results for comparisons of craniocaudal length and mediolateral width between natural and artificial trials and among individuals.**

| Body Dimension | Comparison | *p* | *p_Holm* |
|---|---|---|---|
| Craniocaudal length | Natural vs Artificial (DH1, DH2, DH3) | (0.859, 0.873, 0.877) | - |
| | DH1 vs DH2 | $3.620 \times 10^{-19}$ | $1.085 \times 10^{-18}$ |
| | DH1 vs DH3 | $2.891 \times 10^{-17}$ | $5.770 \times 10^{-17}$ |
| | DH2 vs DH3 | $2.46 \times 10^{-10}$ | $2.463 \times 10^{-10}$ |
| Mediolateral width | Natural vs Artificial (DH1, DH2, DH3) | (0.250, 0.429, 0.695) | - |
| | DH1 vs DH2 | $1.142 \times 10^{-16}$ | $2.284 \times 10^{-16}$ |
| | DH1 vs DH3 | 0.002 | 0.002 |
| | DH2 vs DH3 | $1.737 \times 10^{-17}$ | $5.212 \times 10^{-17}$ |

Holm-Bonferroni corrected *p*-value (*p_Holm*).

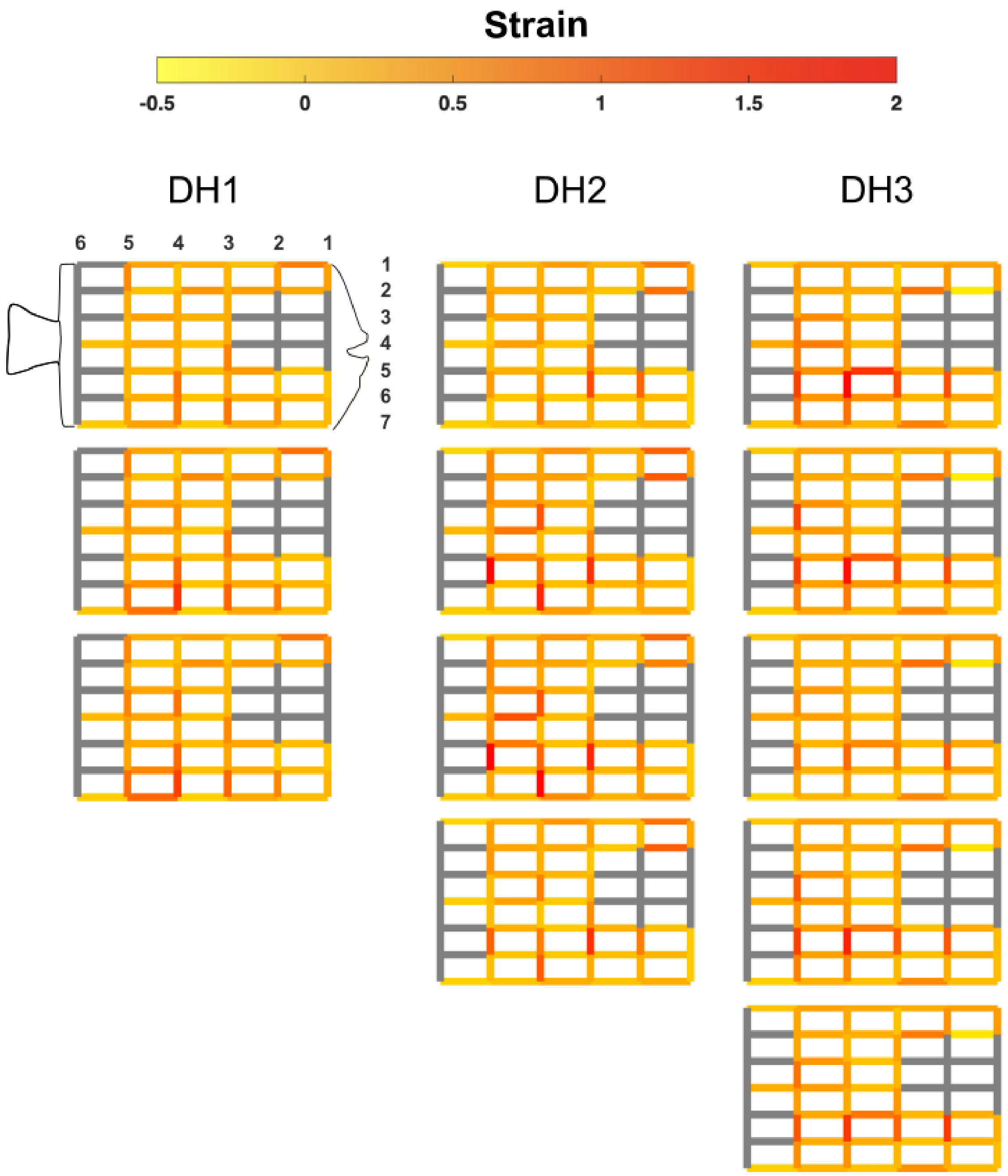


**Figure S2. Colourmap of all axial and hoop skin strains during natural inflation trials.** Skin strains observed in the axial (rows) and hoop (columns) direction at maximum inflation. Each column of colourmaps are the trials for each fish (DH1 *n* = 3, DH2 *n* = 4 and DH3 *n* = 5, 12 natural trials in total from 3 individuals). All colourmaps/grids follow the same format shown in row 1 column 1 where rows/columns are numbered and cranial is pointing right and caudal is left. All grey lines are locations where no skin strain was calculated due to anatomical landmark obstruction e.g., the eye and pectoral fin.

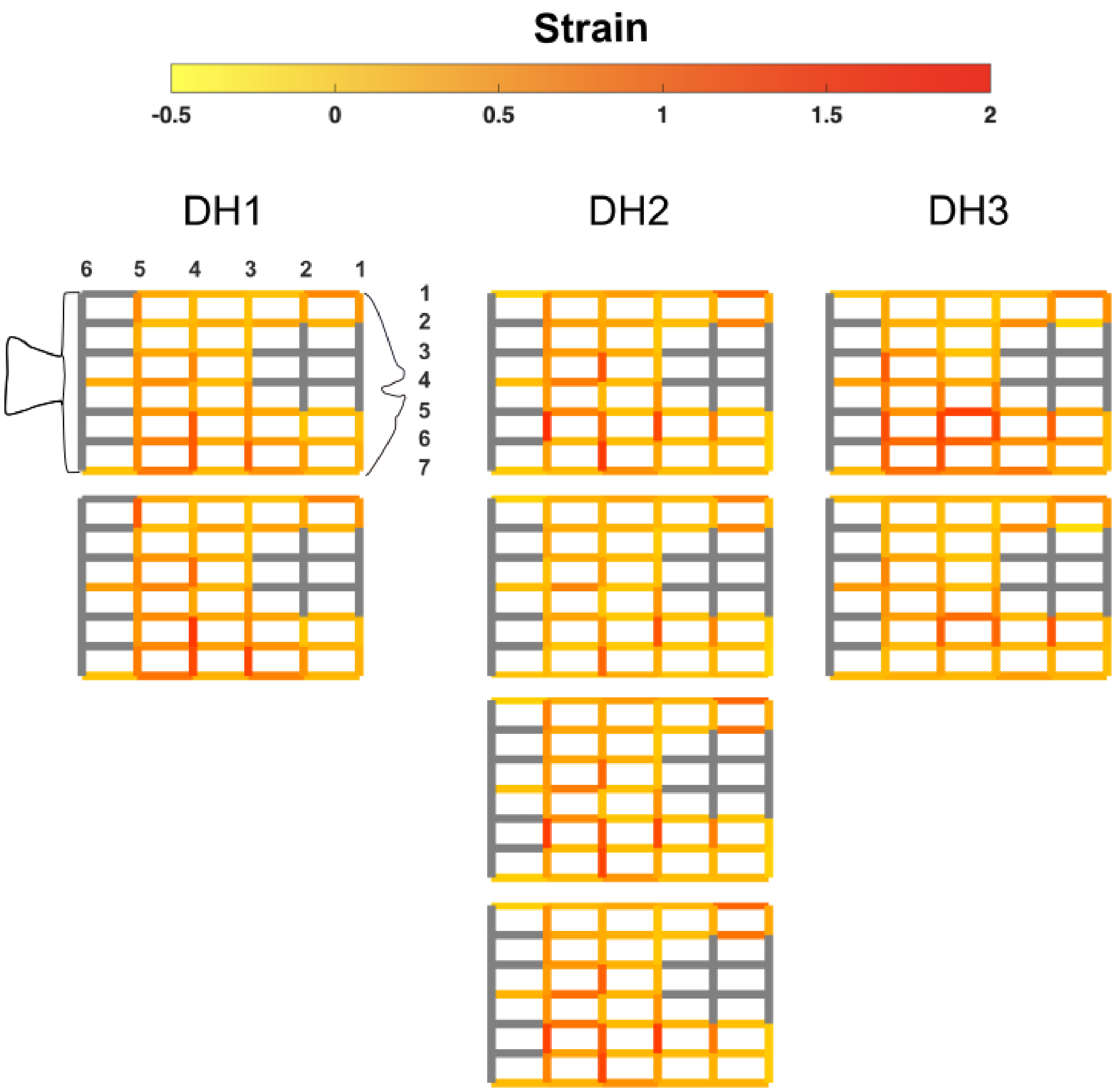


**Figure S3. Colourmap of all axial and hoop skin strains during artificial inflation trials.** Skin strains observed in the axial (rows) and hoop (columns) direction at maximum inflation. Each column of colourmaps are the trials for each fish (DH1 *n* = 2, DH2 *n* = 4 and DH3 *n* = 2, 8 natural trials in total from 3 individuals). All colourmaps/grids follow the same format shown in row 1 column 1 where rows/columns are numbered and cranial is pointing right and caudal is left. All grey lines are locations where no skin strain was calculated due to anatomical landmark obstruction e.g., the eye and pectoral fin.
